# Coexistence of asymmetric cell shape dynamics governs organisational diversity in sensory epithelia

**DOI:** 10.1101/2025.11.03.686357

**Authors:** Anubhav Prakash, Raman Kaushik, Nishant Singh, Ankita Walvekar, Sradha Saji, Raj K Ladher

**Author notes:** **Authors for Correspondence:** Raj K. Ladher, Anubhav Prakash. Max-Planck Institute for Molecular Bio-Medicine, Röntgenstraße 20, Münster, Germany, 48149 (Anubhav Prakash).

## Abstract

Tissues and organs develop a wide range of shapes, sizes and cellular organisations, linked to their physiological function. How developmental programmes generate this organisational diversity remains poorly understood. Inner ear sensory epithelia in birds, fish, and mammals provide a tractable system to address this question, as their constituent cells arise from a common developmental lineage, share molecular signatures, and have conserved physiological functions, yet assemble into distinct cellular organisations. These epithelia contain two principal cell types, mechanosensory hair cells (HCs) and supporting cells (SCs), organised into mosaics. Here, we developed a three-parameter morphospace that quantitatively captures epithelial organisation, enabling comparisons across species, structures and developmental stages. Combining this framework with genetic perturbation, biochemical analysis and live imaging reveals that the diverse mature organisations emerge from a common early organisational state constrained by Notch-Delta signalling. During subsequent development, the differential localisation of adhesion molecules (Cdh1,2 and Nectin) and contractility-associated proteins (NM2 and α-actinin-4) at HC-SC and SC-SC junctions establishes junctional asymmetries. Across species, asymmetry drives stable circular HCs and actively remodelling SCs. The coexistence of these cell shape states drives selective intercalation, guiding each epithelium along a distinct trajectory through morphospace toward its mature organisation. These findings identify cell-shape dynamics as a developmental mechanism for generating organisational diversity in sensory, and potentially other, epithelia.

## Introduction

Developmental programmes integrate genetically encoded information and mechano-chemical feedback to generate the diverse shapes, sizes and cellular organisation of tissues and organs (Collinet and Lecuit, 2021; Bailles, Gehrels and Lecuit, 2022; Rombouts, Elliott and Erzberger, 2023). This diversity can be represented as a morphospace, a multi-dimensional phenotypic space in which each axis captures a defining feature of tissue organisation (Pie and Weitz, 2005; Budd, 2021). Mapping how developing tissues navigate through this morphospace and occupy specific regions provides a route to understanding how developmental mechanisms generate organisational diversity.

Inner ear sensory epithelia are well-suited to this problem. In vertebrates, the inner ear arises from the otic placode. A combination of WNT, BMP, FGF and SHH signalling specifies a pro-sensory domain (Driver *et al*., 2008; Groves and Fekete, 2012; Mann *et al*., 2014; Ono *et al*., 2014; Thiede *et al*., 2014; Huh, Warchol and Ornitz, 2015; Basch *et al*., 2016) that gives rise to at least 6 sensory structures: three cristae, the utricle, the saccule, and an auditory epithelium (Mann *et al*., 2017) (Fig. 1A). Each sensory epithelium consists of two cell types: mechanosensory hair cells (HCs) and supporting cells (SCs), whose fates are specified through Notch-Delta lateral inhibition (Bermingham *et al*., 1999; Lanford *et al*., 1999) (Fig.1B). Despite this conserved developmental logic and shared molecular identities, mature sensory epithelia differ in their cellular organisations across species and structures. This combination of conservation and diversity makes the inner ear an exceptional system to investigate how common developmental programmes generate divergent tissue architecture.

**Figure 1:**
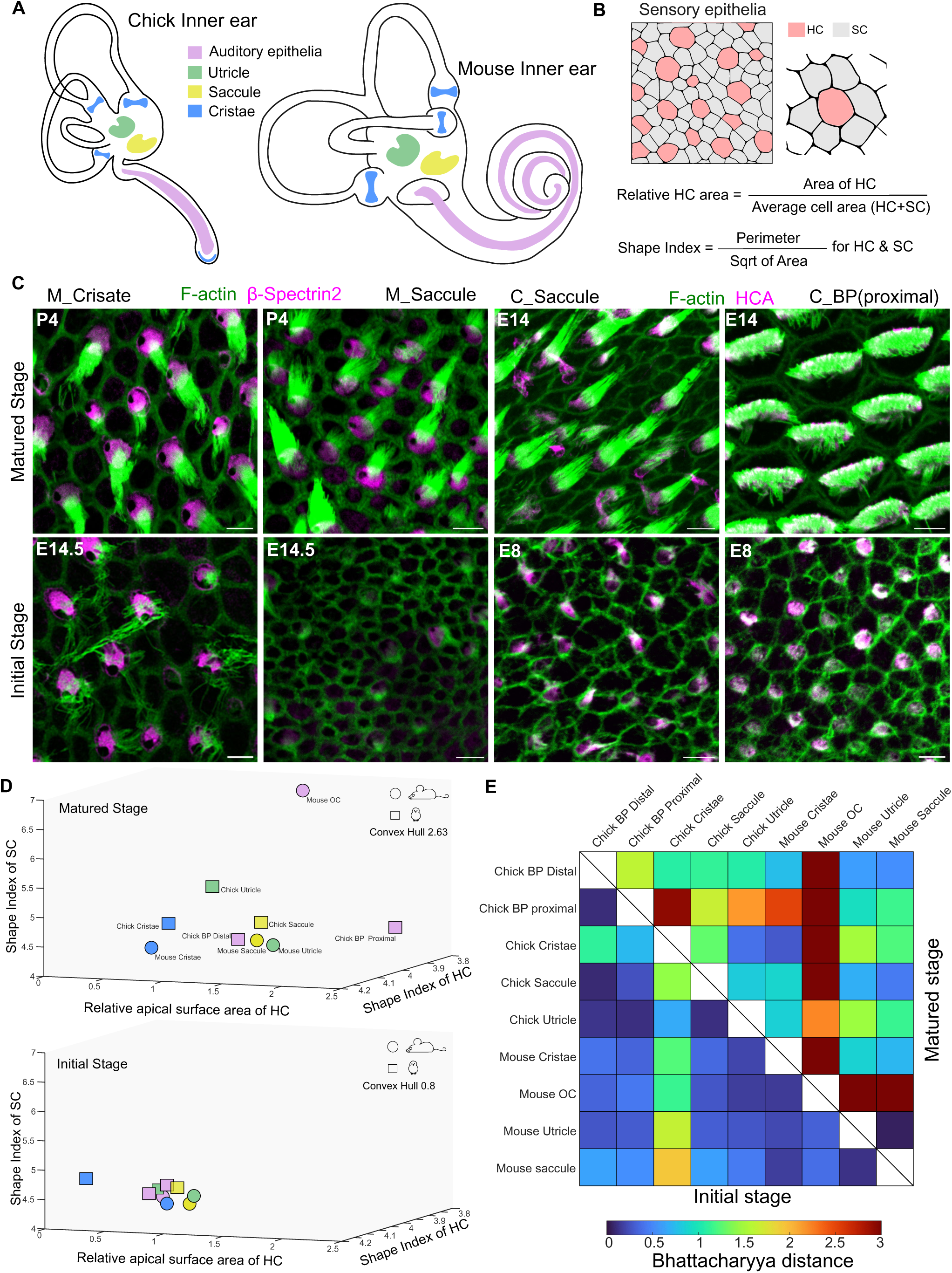
Diverse sensory epithelial organizations emerge from a common initial morphospace. A. Schematic of avian and mammalian inner ear representing the auditory and vestibular sensory structures. B. Schematic of notch-delta signalling drives differentiation of HCs and SCs from the pro-sensory domain and the calculation used for morphospace calculations. C. Sensory epithelia of mouse Cristae, saccule and chick BP and saccule at the matured stage (P4 for mouse and E14 for chick) and initial stage (E14.5 for mouse and E8 for chick) stained for F-actin (green) and hair cell marker (HCA, Beta Spectrin 2). D. A 3-dimensional morphospace representing the HC-SC mosaic for relative apical surface area, shape index of HC and SC at the matured stage and initial stage. N= 4 samples each for mouse and chick sensory epithelia. E. Bhattacharyya distance that measures the overlap between the groups for initial and mature stages. Scale Bar: 5µm See Figure S1 and S2 for details

In epithelia, cellular organisation reflects the shape and neighbour relationships of cells. These properties arise from the balance between junctional adhesion, mediated by molecules such as cadherins and nectins, and actomyosin-driven contractility (Duguay, Foty and Steinberg, 2003; Togashi *et al*., 2011; Wickström and Niessen, 2018; Cohen *et al*., 2020; Tsai *et al*., 2020; Prakash, Raman, *et al*., 2025; Prakash, Weninger, *et al*., 2025). Two-dimensional vertex models have formalised this principle in simple epithelia and used ground-state diagrams to describe organisational diversity in single-cell-type epithelia (Farhadifar *et al*., 2007; Muthukrishnan *et al*., 2026). In such tissues, integrated features of cell shape are shown to transition with tissue-scale organisation, allowing cross-species comparison (Andrews *et al*., 2021; Mendieta-Serrano *et al*., 2025; Battistara *et al*., 2026). However, a quantitative framework that explains the organisation of epithelia with multiple cell types and is implemented for cross-comparison is missing.

Here, we develop a minimal three-parameter morphospace for vertebrate epithelia and use it to compare organisation across species, structures and time. We show that sensory epithelia begin from an initial Notch-Delta-constrained organisational state and subsequently diverge through tissue-specific trajectories. This divergence is driven by differential localisation of junctional adhesion and contractility components, which produce stable circular HCs and dynamically remodelling SCs. Together, the coexistence of this asymmetric cell-shape dynamics governs directed intercalation and guides each epithelium toward its mature organisation.

## Results

### Diverse sensory epithelia organisation emerges from a common initial state

To investigate organisational diversity across sensory epithelia, we analysed stage-fixed mouse and chick inner ears. We first examined mature stages, when final cellular patterns are established: embryonic day 14 (E14) in the chick and postnatal day 0 (P0) in the mouse for the auditory epithelia and postnatal day 4 (P4) for the mouse vestibular epithelia(McKenzie, Krupin and Kelley, 2004; Burns *et al*., 2012, 2015). Junctional actin was labelled with phalloidin to segment cell boundaries, and HCs were identified using hair cell antigen (HCA) in chick and β-spectrin II in mouse.

Across sensory structures, HCs and SCs shared a common mosaic arrangement, with HCs interspersed with SCs. However, the apical surface area of both cell types varied across epithelia, ranging from approximately 10µm^2^ −100µm^2^ for HC and 2µm^2^ - 50µm^2^ for SCs (Fig. 1C, S1A-C). To compare these epithelia quantitatively, we mapped two phenotypic descriptors commonly used for simple epithelia onto a three-axis morphospace. The first axis was relative apical surface area, defined as the area of HCs divided by the average cell area in the epithelium (Fig. S1D-E). The second and third axes were the shape indices of HCs and SCs, calculated as perimeter divided by the square root of area (Fig. 1B). Mature sensory epithelia occupied distinct regions in this morphospace (Fig. 1D). Vestibular epithelia clustered relatively closely across species, whereas auditory epithelia occupied more distant regions. Pairwise two-dimensional projections confirmed that vestibular epithelia were similar but not identical (Fig. S2A-C). To quantify separation between epithelial distributions, we used Bhattacharyya Distance (BD), in which low BD values indicate substantial overlap, while high values indicate well-separated distributions with minimal overlap (Kailath, 1967). Except for the mouse utricle and saccule, all pairwise comparisons between mature epithelia had BD values greater than 1 (Fig. 1E). Thus, mature sensory epithelia are quantitatively distinct and separable in morphospace.

The diversity observed at maturity could arise from differences in initial cellular patterning, developmental remodelling, or from both. To assess the contribution of initial cellular patterning, we immunostained sensory epithelia at the earliest stage when HCs are first apparent: E8 in chick and E14.5-15.5 in mouse. At this stage, HCs from all sensory structures displayed apical surface areas smaller than SCs (Fig.1C; S1A, D-E). Strikingly, these sensory epithelia occupied a small, overlapping region of morphospace (Fig. 1D). The total morphospace volume decreased from 2.63 at maturity to 0.8 at the initial stage. Pairwise projections and reduced BD values further confirmed the high overlap among early-stage epithelia (Fig. 1E, S2A-C). These data suggest that sensory epithelia begin development with limited organisational diversity and diverge towards distinct mature architectures.

### Lateral Inhibition-based fate specification constrains initial geometry

We next asked what underlies the conserved early organisation of HCs and SCs. The role of Notch signalling in HC-SC differentiation has been well-established across vertebrate sensory epithelia (Lanford *et al*., 1999). Here, lateral inhibition acts on an initially equivalent field into Delta-high cells that become HCs and Notch-high cells that become SCs (Fig. 2A, B). Theoretical models and synthetic biology studies of Notch-Delta signalling further predict that cells with smaller apical surface areas preferentially adopt a Delta-high state (Jolly *et al*., 2015; Hunter *et al*., 2016; Shaya *et al*., 2017; Kasirer and Sprinzak, 2024; Singh *et al*., 2024). Our finding that nascent HCs are smaller and circular (Fig. 1C; S1A, D, E) suggested that Notch-Delta lateral inhibition might also impose the conserved geometric bias.

**Figure 2:**
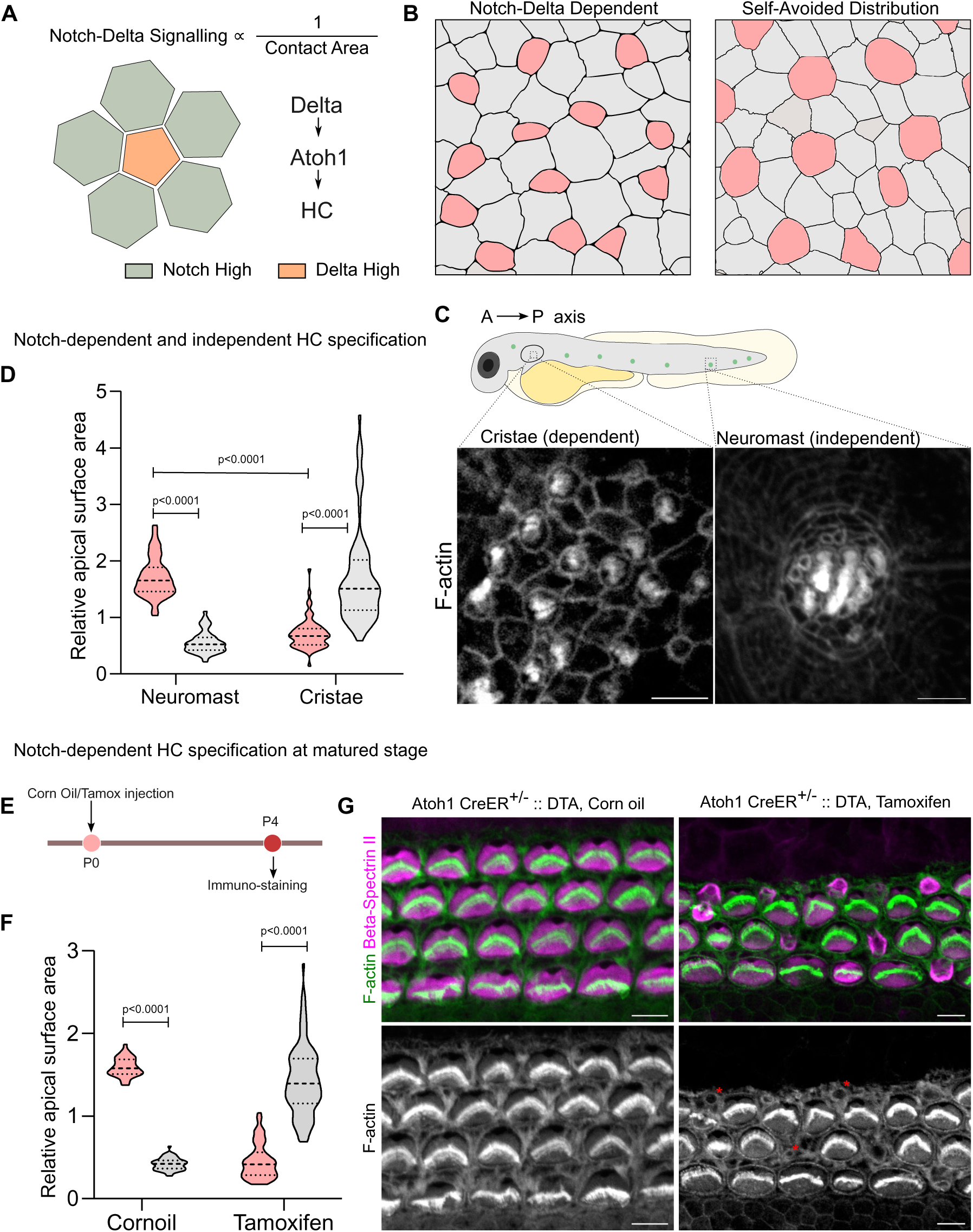
Lateral inhibition-based fate specification constrains initial geometry. A. Schematic showing the relation between cell contact area and Notch-Delta signalling. B. Notch-Delta dependent and independent self-avoided distribution mechanisms for initial HC-SC organisation in sensory epithelia C. 3 days post fertilisation zebrafish cristae and neuromast stained for F-actin (grey), showing HC-SC mosaic. N= 15 cristae and 28 neuromast from 12 embryos. D. Relative apical surface area of HCs and SCs for cristae and neuromast. N= 72/98 neuromast, 67/89 cristae (HC/SC). E. Schematic showing the experimental setup for HC loss and recovery. F. Relative apical surface area of HCs and SCs from OC of corn oil injected and tamoxifen-induced Atoh1 CreER^+/-^ :: DTA embryos at P4. N= 45/54 corn oil, 42/48 tamoxifen (HC/SC). G. P4 OC from corn oil injected and tamoxifen induced Atoh1 CreER^+/-^ :: DTA embryos stained for F-actin (green and grey) and beta-spectrin II (magenta). Red asterisks show neonatal HCs. N= 4 embryos H. Relative apical surface area of HCs and SCs from sensory epithelia of chick. N= 65/138 BP, 64/148 Utricle, 61/138 Saccule, 78/152 cristae (HC/SC). Scale Bar: 5µm Statistics: Unpaired Test

To test this, we used a natural comparison between two sensory structures of zebrafish. Zebrafish offer a direct test of this idea, because their sensory structures build HCs by distinct cellular routes. In inner ear sensory structures such as the cristae, HC–SC fate is resolved by Notch-Delta lateral inhibition acting across a pro-sensory field, as is the case in the mouse and chick inner ear (Bermingham *et al*., 1999; Lanford *et al*., 1999). In the neuromasts of the lateral line, Notch-Delta signalling specifies the progenitor of the HC, which, through a terminal, symmetric division, generates pairs of sibling HCs. SCs are maintained as a separate, self-renewing population (Puligilla *et al*., 2007; Wibowo *et al*., 2011; Lush *et al*., 2019). This difference allows us to ask whether the conserved geometry is specifically tied to lateral-inhibition-based specification (Fig. 2B). At three days post-fertilisation, when HCs first become apparent, cristae HCs were smaller and more circular than surrounding SCs. In contrast, neuromast HCs were significantly larger than SCs (Fig. 2C, D), consistent with lateral inhibition biasing the geometry of the sensory epithelium.

The early postnatal mammalian cochlea retains a transient capacity to regenerate. Following HC loss, Notch signalling from the lost HC is withdrawn, and lateral inhibition is released. In a process that recapitulates differentiation, a subset of SCs de-repress Atoh1 and transdifferentiate into new HCs (Burns and Stone, 2017; Li *et al*., 2025).We reasoned that if the small, circular geometry of nascent HCs is a signature of lateral-inhibition-based specification rather than of embryonic developmental stage, then a HC newly generated by this mechanism amongst mature HCs should adopt the same geometry. To test this, we used a tamoxifen-inducible, HC-specific driver (Atoh1-CreERT2) to express diphtheria toxin fragment A (DTA) in early postnatal HCs. Following tamoxifen induction at birth, new HCs, expressing by β-spectrin II and the absence of stereocilia, were observed at P4 (Fig. 2E), but not in corn-oil-injected controls. These nascent HCs were significantly smaller and more circular than their neighbouring SCs (Fig. 2F, G). This suggests that when lateral inhibition operates within a mature cochlear epithelium, it again produces small, circular HCs, indicating that this geometry reflects the mode of HC specification rather than the developmental state of the tissue.

Together, these findings suggest that Notch-Delta signalling imposes a conserved geometric bias across vertebrate sensory epithelia, constraining their initial organisation to a common, overlapping region of morphospace.

### Epithelia diverge through selective neighbour exchange

We next asked how developmental remodelling contributes to the diversity in organisation at the mature stage. To address this, we tracked morphospace trajectories across developmental stages in three epithelia representing the full range of mature organisational outcomes: The mouse cristae, where SCs are larger than HCs at maturity; mouse saccule, where HCs and SCs reach comparable apical surface areas; and chick basilar papilla, where HCs become larger than SCs (Fig. 3A, B). Despite their distinct mature organisations, all three epithelia initially underwent a relative increase in HC apical surface area, converging on a shared intermediate organisational state in which HC and SC apical surface areas are comparable (Fig. 3A, B). This intermediate was reached at different absolute developmental timepoints, E10 for BP, E16.5 for saccule and E18.5 for cristae, after which each epithelium followed a distinct trajectory through morphospace toward its mature organisation. Since cell proliferation is largely complete by this stage, we hypothesised that divergence is driven by regulated neighbour exchange rather than by continued growth of the population (Goodyear and Richardson, 1997; Burns *et al*., 2012; Yang *et al*., 2017; Prakash, Weninger, *et al*., 2025). To test this, we quantified the change in neighbour number for HCs and SCs across development. In BP, mean HC neighbour number increased from an average of 4 to 8, while SC neighbour number decreased from 5 to 4. In saccule and cristae, both HC and SC neighbour numbers remained closer to 5 throughout development (Fig. 3C). Across all three epithelia, the number of SC neighbours shifted during development with SCs progressively losing SC-SC contacts and gaining HC-SC contacts. To quantify this, we measured the fraction of HC–SC junctions across developmental stages. In BP, the fraction of HC–SC junctions increased from 20% to 40%. In saccule, it rose from 18% to 27%, and in cristae from 22% to 27%. The extent of change in HC–SC junction fraction scaled with the distance each epithelium travelled through morphospace to reach maturity.

**Figure 3:**
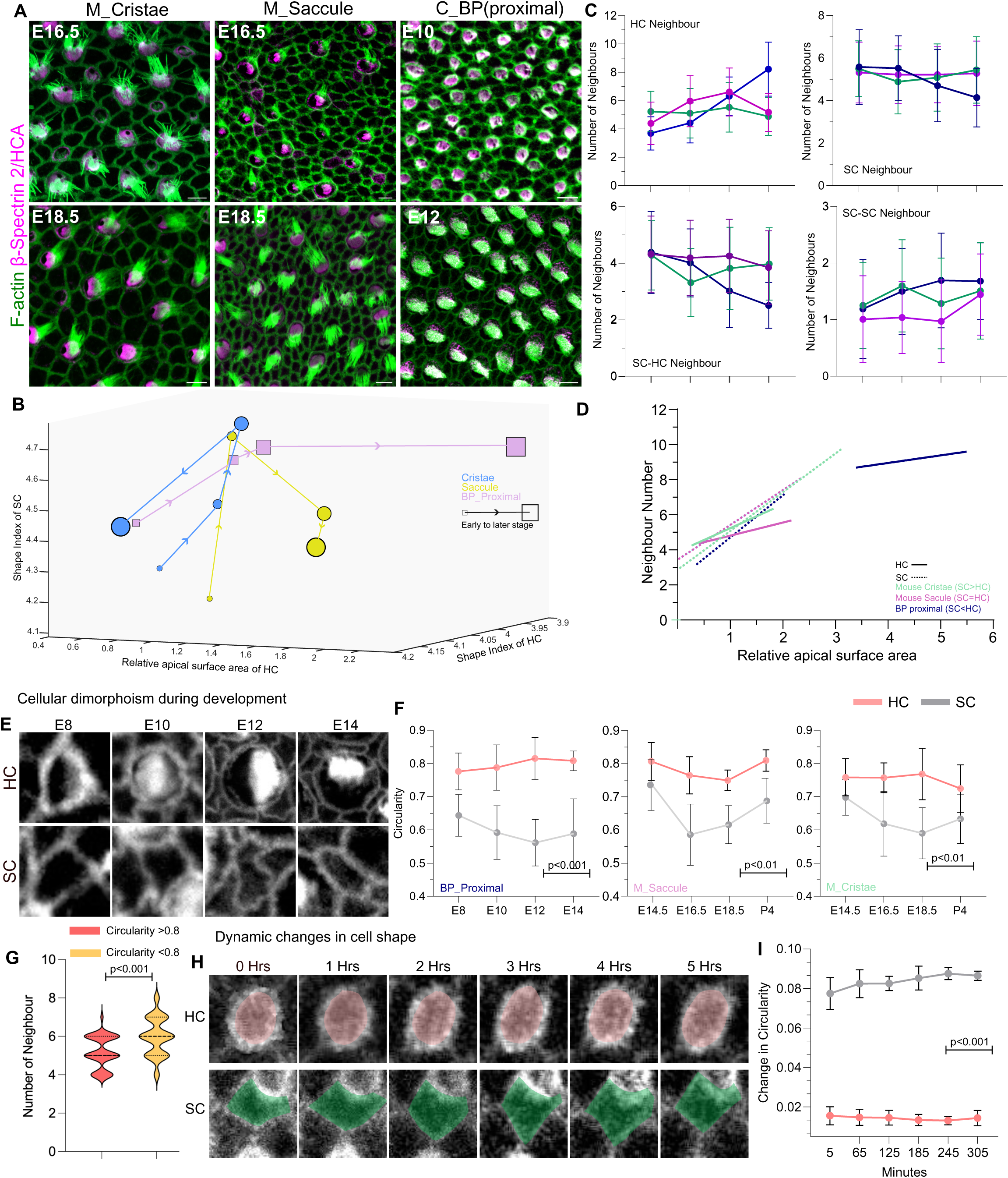
Cellular dimorphism regulates neighbour exchanges and governs epithelial organisation. A. Mouse cristae, saccule and BP stained for F-actin(green) and hair cell marker, Beta spectrin 2 and HCA (magenta) at E16.5 and E18.5 in mouse and E10 and E12 in chick. B. Morphospace trajectories for mouse cristae(blue), saccule(yellow) and chick BP (purple) from initial (small symbol) to final stages (large symbol). C. Number of neighbours for HCs and SCs for mouse cristae (green), saccule (magenta) and proximal part of BP (blue) during development from initial to final stage. N=3 samples for each tissue each stage. D. Linear regression graph representing the relation between the relative apical surface area and number of neighbours for HCs (bold lines) and SCs (dashed line) in mouse Cristae, saccule and chick BP proximal. E. Representative image of HCs and SCs stained for F-actin from E8 to E14, representing the generation (E8-10) and maintenance (E10-14) of compact shape for HC and elongated shape for SC in BP. F. Circularity of HC (pink) and SC (grey) between E8-E14 for chick BP, mouse saccule and mouse cristae. G. Number of neighbours for cells from E10 BP with circularity index higher or lower than 0.8. H. Snapshots from live imaging of E10 BP at 0, 1,2,3,4, 5 hours showing compact shape for HCs (pink shading) and dynamic changes in SCs (green shading) shape. I. Change in circularity for HC (pink) and SC (grey) in 5 hours, calculated every 1 hour.N=3 videos/ 40 cells for HCs and SCs. Scale Bar: 5µm Statistics: Unpaired-Test, Bars represent std. Deviation.

Together, these data show that SCs selectively exchange SC neighbours for HC neighbours during development, and that the extent of this neighbour exchange corresponds to each epithelium’s final position in morphospace.

### Asymmetric cell shape dynamics govern selective intercalation

To understand the basis of this directed neighbour exchange, we turned to Lewis’ law, the empirical linear relation between cells apical surface area and its number of neighbours (Lewis, 1926; Kokic *et al*., 2019). Both HCs and SCs obeyed this relationship, with neighbour number rising with apical area. Crucially, however, HCs showed a consistently shallower slope than SCs across all three epithelia (Fig. 3D; Fig. S3): at any given surface area, an HC had fewer neighbours than an SC. Apical area alone therefore cannot account for the directed intercalation. Reasoning that cell shape might supply the missing variable, we found that HCs across these structures attained a circular shape significantly different then SC during development (Fig. 3E-F). For both HCs and SCs, at equal surface area cells of higher circularity had fewer neighbours (Fig. 3G). This suggested that cell shape, not just size, explains the neighbour composition.

If cell shape sets neighbour number, then differences in shape dynamics between HCs and SCs should determine which junctions are remodelled, and therefore the direction of neighbour exchange (Curran *et al*., 2017; Wang *et al*., 2020). We tested this by live-imaging shape change directly. We stained E10 BP, the intermediate stage, with SiR-actin to label cell boundaries and allowing identification of HCs by labelling stereociliary bundles and imaged for 5 hours (Movie 1). Over this window, HCs underwent significantly smaller changes in circularity than their SC neighbours, such that within a single epithelium, HCs held a stable, compact shape with change in circularity at 0.01± 0.001 every hour while SCs remodelled continuously with change in circularity at 0.08 ± 0.003 in every hour (Fig. 3H-I). Thus, within a single epithelium, HCs and SCs significantly (P<0.001) differ in their shape dynamics, with HCs maintaining a stable circular shape while SCs remodel.

As cell shape are governed by adhesion and contractility at cellular interfaces, we next asked whether HC–SC and SC–SC junctions differ in composition (Fig 4A). Cdh2, the dominant cadherin in BP, and Nectin1 were enriched at HC–SC junctions relative to SC–SC junctions (Fig. 4B-E). The actin cross-linker Actn4, which couples the junctional complex to the actomyosin network, was similarly enriched at HC–SC junctions, where it co-localised with non-muscle myosin IIB and the phosphorylated form of the regulatory light chain (RLC) of non-muscle myosin II (Fig. 4G-I). Thus, HC–SC and SC–SC junctions carry distinct complements of adhesive and contractility proteins. To test whether this asymmetry underlies the difference in shape dynamics and neighbour exchange, we perturbed junctional contractility and Actn4 directly. Culturing BP with the Rho-kinase inhibitor Y-27632, lowered non-muscle myosin activity. We find that it also reduced the asymmetry of Actn4, reduced SC shape change, increased HC shape change, and reduced the difference of neighbour numbers between HC and SC (Fig. 4J-N, Movie 2). We then disrupted Actn4 specifically by CRISPR-Cas9 mosaic knockout, electroporating the E3.5 avian otic primordium with a construct co-expressing GFP and either a control-gRNA, or Actn4-gRNA, and analysing GFP-positive patches at E10 (Singh *et al*., 2022). Control patches showed Actn4 localisation comparable to surrounding tissue, whereas Actn4-gRNA-targetted patches showed markedly reduced Actn4 (Fig. S4). HCs within these knockout patches showed reduced circularity and increased neighbour number (Fig. 4O-Q).

**Figure 4.**
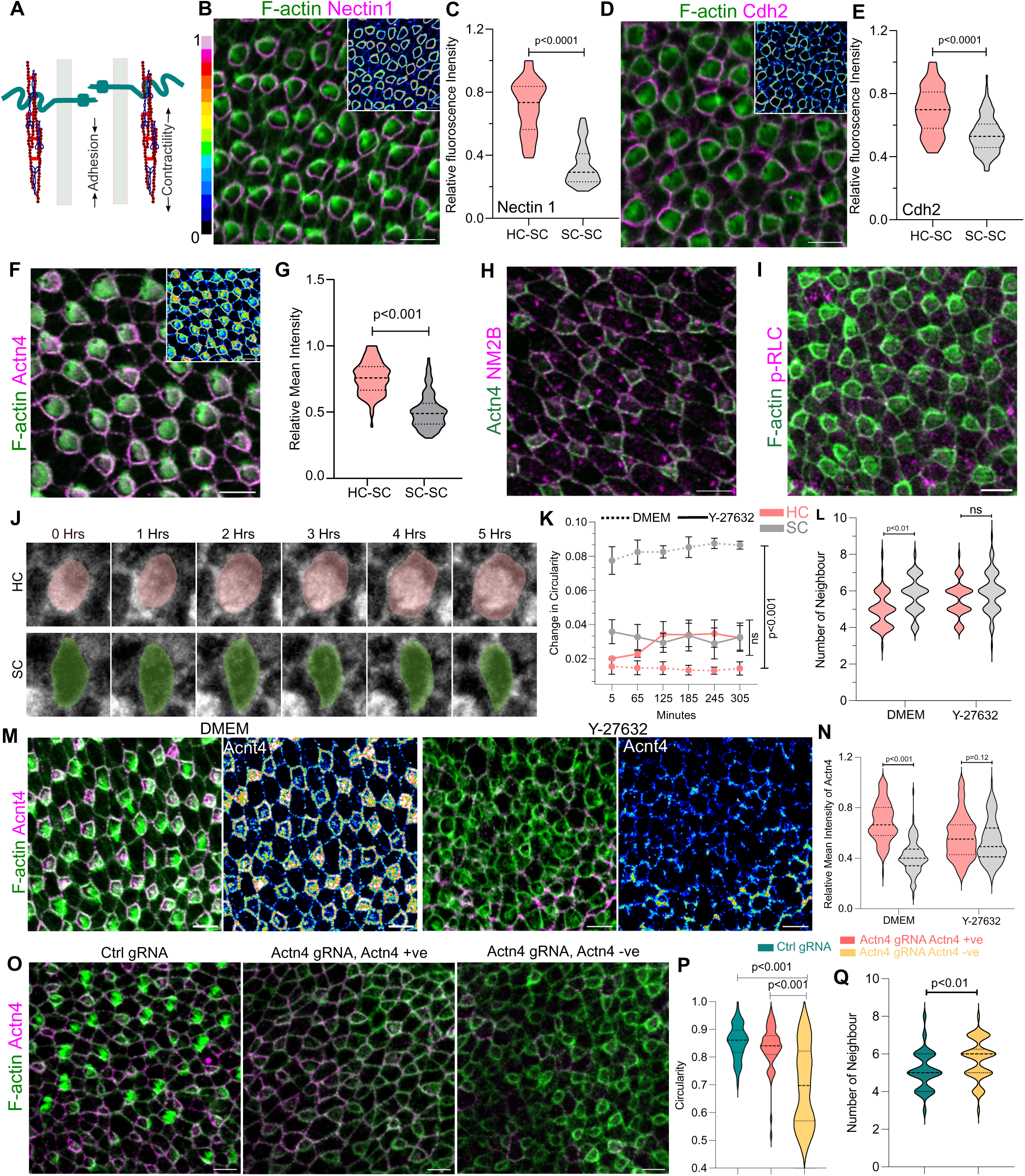
Junctional heterogeneity governs cellular dimorphism. A. Schematic representing the role of adhesion and contractility in cell-cell interface. B. E10 BP stained for F-actin (green) and Nectin-1 (Magenta,16colors LUT). C. Relative Mean intensity of Nectin 1 on HC-SC and SC-SC junctions. D. E10 BP stained for F-actin (green) and Cdh2 (Magenta, 16colors LUT). E. Relative Mean intensity of Cdh2 on HC-SC and SC-SC junctions. F. E10 BP stained with F-actin (green) and Actn4 (Magenta, 16colors LUT). G. Relative Mean intensity of Actn4 on HC-SC and SC-SC junctions. H. E10 BP stained with Actn4 (green) and NM2B (Magenta). I. E10 BP stained with F-actin (green) and active form of NM2B (phosphorylated-RLC) (Magenta). J. Snapshots from live imaging of E10 BP at 0, 1,2,3,4, 5 hours showing change in shape for HCs (pink shading) and SCs (green shading). K. Change in circularity for HC and SC after inhibition of Myosin activity by 10µM Y-27632 compared to the control DMEM. N=3 videos/40 cells L. Number of neighbours for HCs and SCs for control E10 BP grown in explants for 4 hours and E10 BP grown in presence of Y-27632 (10µM) for 4 hours. M. E10 BP explant for 4 hours in control (DMEM) and Y-27632 (10µM) stained for F-ctin (green) and Actn4 (magenta, 16 color LUT). N. Relative Mean intensity of Actn4 on HC-SC and SC-SC junctions in DMEM and Y-27632 treated BP. O. E10 BP electroporated with ctrl gRNA, Actn4 gRNA but positive for Actn4 staining (non-electroporated patches) and Actn4 gRNA negative for Actn4 staining (Actn4 KO patches) stained for F-actin (green) and Actn4 (magenta). N=4 samples P. Circularity of HCs from E10 BP electroporated with ctrl gRNA, Actn4 gRNA but positive for Actn4 staining (non-electroporated patches) and Actn4 gRNA negative for Actn4 staining (Actn4 KO patches). N=4 samples each and 78/87/68 cells. Q. Number of neighbours for HCs from E10 BP electroporated with ctrl gRNA, Actn4 gRNA but positive for Actn4 staining (non-electroporated patches) and Actn4 gRNA negative for Actn4 staining (Actn4 KO patches). N=4 samples each and 78/87/68 cells. Scale Bar: 5µm Statistics: Unpaired-Test, ns= P>0.05

Together, these data suggest that HC–SC and SC–SC junctions differ in the enrichment of Cdh2, Nectin1, Actn4, and active myosin IIB, that HCs and SCs differ in apical shape dynamics, and that reducing junctional contractility or Actn4 abolishes both the shape-dynamics asymmetry and the directed difference in neighbour number.

### Junctional asymmetry shapes organisational trajectories

We next asked whether the junctional asymmetry that drives selective intercalation also governs organisational divergence across epithelia. We first established that this asymmetry is a shared feature of the epithelia we compare. In cristae, both Cdh1 and NM2A were enriched on HC-SC junctions when compared to SC-SC junctions (Fig. 5A). In saccule Cdh1 was enriched on SC-SC junctions and NM2B was enriched on HC-SC junctions (Fig. 5B). This data show junctional asymmetry is therefore a conserved feature across sensory epithelia spanning distinct organisational classes while the molecules defining that heterogeneity may vary.

**Figure 5.**
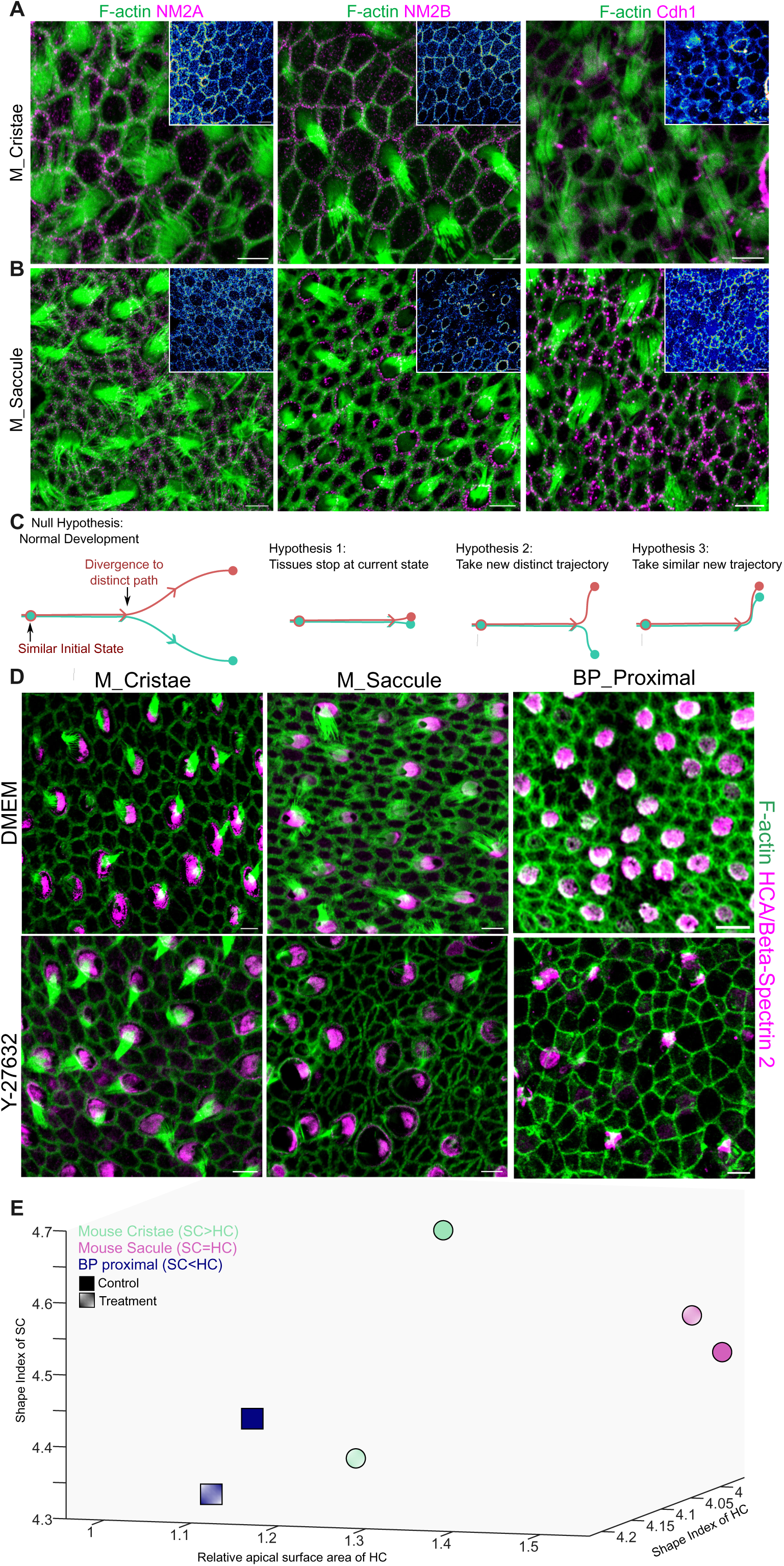
Junctional heterogeneity guides epithelia to distinct morphospace trajectory. A. E18.5 mouse cristae stained for F-actin (green), NM2A (magenta, 16 colours LUT), NM2B (magenta, 16 colours LUT) and Cdh1 (Magenta, 16 Colours LUT). B. E18.5 mouse saccule stained for F-actin (green), NM2A (magenta, 16 colours LUT), NM2B (magenta, 16 colours LUT) and Cdh1 (Magenta, 16 Colours LUT). C. Hypothesis for how inhibition of junctional heterogeneity can impair the morphospace trajectory. D. Mouse cristae (E18.5), mouse saccule (E16.5) and chick BP (E10) cultured for 12 hours in presence of control (DMEM) and ROCK-inhibitor (Y-27632, 10µM) for 12 hours and stained for F-actin (green) and hair cell marker, Beta Spectrin 2/ HCA (magenta). E. Morphospace showing control (DMEM) and Y-27632 (10µM) treated epithelia occupy distinct regions of morphospace.

Given this shared starting point, three outcomes could follow from removing junctional asymmetry, each making a distinct prediction (Fig. 5C). If the asymmetry is required to maintain organisation, its loss should arrest each epithelium in its current state. If it constrains organisation, its loss should release each epithelium into a new state, distinct both from its own control and from the others. If the asymmetry actively drives divergence, its loss should collapse all three epithelia toward a common state. To test these, we cultured E18.5 mouse cristae, E16.5 mouse saccule, and E10 chick BP representing their intermediate stage (Fig 3A,B) for 12 hours in Y-27632. All three treated epithelia adopted organisations significantly different from their respective controls, ruling out arrest (Fig. 5D,E; Fig. S6A–D). However, the three remained organisationally distinct from one another, ruling out convergence to a common state (Fig. 5D,E; Fig. S6D). Loss of junctional asymmetry thus neither freezes each epithelium nor collapses them together; instead, it redirects each along its own distinct organisational morphospace trajectory.

## Discussion

Our findings describe a conserved developmental logic underlying the diversity of hair-cell (HC) and supporting-cell (SC) organisation across vertebrate sensory epithelia. Each epithelium begins from a common initial geometry, starting from small, circular HCs among larger SCs yet reaching a distinct mature organisation, from SCs larger than HCs in the cristae to HCs larger than SCs in the chick basilar papilla (BP). Despite these different endpoints, the developmental trajectories of all three epithelia we compared, mouse cristae, mouse saccule, and BP, converge on a shared intermediate state in which HC and SC apical areas become comparable, at markedly different absolute times (E10 in BP, E16.5 in the saccule, E18.5 in the cristae), indicating a conserved developmental logic rather than synchronised timing. We propose this intermediate stage as an attractor state, from which each epithelium then resolves towards its own mature identity.

The initial small, circular HC geometry is a signature of lateral-inhibition-based specification (Jolly *et al*., 2015; Shaya *et al*., 2017). This bias emerges wherever HCs are selected from an equivalence group through Notch-Delta lateral inhibition: in the cristae, as in the chick and mouse inner ear more generally, and even in the mature mouse organ of Corti (OC), where inducing HC loss releases neighbouring supporting cells from Notch-mediated inhibition and triggers their transdifferentiation into new HCs (Singh *et al*., 2024; Khalaily *et al*., 2026). These regeneration-derived HCs reactivate the same lateral-inhibition circuit used in development and adopt the same small, circular geometry despite arising within an otherwise mature epithelium, showing that developmental stage cannot account for the bias. By contrast, the bias is absent where HCs arise through a different route, such as the progenitor divisions that generate sibling hair-cell pairs in the zebrafish neuromast (Wibowo *et al*., 2011; Li *et al*., 2025). The conserved geometry is therefore a signature of this specific mode of specification, that is, lateral inhibition, resolving an equivalence group.

From this shared intermediate, each epithelium diverges towards its mature organisation through selective intercalation, in which SCs progressively exchange SC–SC contacts for HC–SC contacts. In BP, HC–SC junctions were enriched for the cadherin Cdh2, the adhesion molecule Nectin1, the actin cross-linker Actn4, and active non-muscle myosin IIB, while SC–SC junctions lacked this composition; a comparable asymmetry in cadherin and myosin distribution was also conserved in the cristae and saccule. We propose that this molecular asymmetry sets the direction of intercalation. Although myosin enrichment marks HC–SC junctions as sites of elevated tension, Actn4-mediated cross-linking may render these junctions resistant to remodelling rather than prone to shrinkage, so that HCs act as stable anchors around which the more compliant, Actn4-poor SC–SC junctions are preferentially resolved during packing (Ehrlicher *et al*., 2015; Kemp and Brieher, 2018). This model is consistent with our finding that HCs maintain a stable, circular shape while SCs continuously remodel within the same epithelium, and with the parallel loss of both the shape-dynamics asymmetry and the neighbour-number asymmetry following disruption of contractility (Y-27632) or of Actn4 itself. Junctional asymmetry therefore acts as a directional guide for remodelling, not a binary determinant of organisational identity.

Selective intercalation driven by asymmetric junctional composition may represent a general mechanism for generating organisational diversity in other mixed epithelia beyond the inner ear, including mucociliary, olfactory, and retinal epithelia, wherever two or more cell types must be patterned across a shared apical surface (Salbreux *et al*., 2012; Katsunuma *et al*., 2016; Kasirer and Sprinzak, 2024; Nommick *et al*., 2025). More broadly, remodelling in these tissues is also shaped by mechanical coupling among junctions, extracellular matrix composition, tissue geometry, and the three-dimensional shape of individual cells (Heisenberg and Bellaïche, 2013; Gnedeva *et al*., 2017; Zaidel-Bar and Agarwal, 2025; Weninger *et al*., 2026). The morphospace framework we present begins to capture these combined inputs as phenotypic outcomes along three quantifiable axes, rather than any single axis, such as relative cell size, in isolation. This allows vertebrate sensory epithelia to be compared directly, and the mechanical, geometric, and biochemical cues that encode their organisational diversity to be dissected. This minimal framework could be extended further, to incorporate three-dimensional phenotypic parameters, tissue-level shape, and orientational features such as planar polarity, providing a more comprehensive basis for comparing epithelial organisation across systems.

## Author Contributions

AP and RKL conceived the project and designed experiments. RKL acquired funding and supervised the project. AP performed experiments and analysed the data. RK designed the Actinin4 gRNA construct and, with NS, performed Actn4 KO experiments. AW helped with explant experiments. SS helped with segmentation of confocal images. AP and RKL wrote the manuscript.

## Supporting information

Code used for analysis

## Acknowledgements

This work was supported by the Department of Atomic Energy, Government of India, Project Identification No. RTI 4006, and grants from SERB, TIFR Infosys-Leading Edge Grant, the Royal National Institute for Deaf People, and the International Foundation for Research and Education via a Simons-Ashoka ECF fellowship to AP, for this research. We thank Central Imaging and Flow cytometry Facility (CIFF), Animal Care and Resource Centre (ACRC) and laboratory kitchen at NCBS. We thank Boby R.V. for help with imaging facility at DBS, Mumbai. We thank CPDO and TI Hasserghatta for providing eggs for experiments. We thank Prof. Mahendra Sonawane, Prof Sandeep Krishna, Dr Tapomoy Bhattacharjee and Prof Hiroshi Hamda for their insightful comments. We thank members of Earlab at NCBS, Bengaluru and members of the Zebrafish epidermis lab at Mumbai for their feedback and help. AP thanks Dr Rajat Mann for his comments on code used for analysis.

## Methods

### Animal Housing

**Mouse.** Mice were housed at animal care and resource centre (ACRC) at National Centre for Biological Sciences (NCBS) in accordance with the approved guidelines by Institutional Animal Ethics Committee (NCBS-IAE-2020/13(R1M_EE)).

**Chick.** Fertilised chicken eggs were procured from Central Poultry Development Organisation & Training Institute, Hasserghatta. Upon arrival at NCBS, it was cleaned and incubated at 37°C and 75% humidity to achieve desired stage.

**Fish.** Zebrafish were housed and handled at Zebrafish facility at Tata Institute of Fundamental Research, Mumbai in accordance with the guidelines recommended by the Committee for the Purpose of Control and Supervision of Experiments on Animals (CPCSEA), Government of India, and approved by the Institutional Animal Ethics Committee (TIFR/IAEC/2024-5).

### Immuno-staining

Mouse embryos were collected from timed pregnant females identified by vaginal plug and Theiler staging series. Chick embryos at specific stages were obtained by cracking open incubated fertilized eggs. The embryos were dissected in ice-cold Phosphate Buffer Saline (without Ca2+ and Mg2+) and the inner ear was fixed in 4% paraformaldehyde (PFA) for 4 hours at room temperature. For fish, transgenic Tg(CldB:LynEGFP) lines were used to collect 3-day-old embryos with marked junctions, which were fixed in 4% PFA for 4 hours at RT.

Dissected sensory epithelia or entire fish (for neuromasts) were permeabilized with 0.3% PBST (Tween-20) for 30 minutes at RT on a shaker. Next, samples were incubated with blocking solution containing 5% heat-inactivated goat serum and 1% BSA in PBST for 1 hour at RT on a shaker. Subsequently, samples were incubated with primary antibodies (HCA at 1:2000; Beta Spectrin II at 1:400; Alpha-actinin 4 at 1:400; NM2B at 1:400) overnight at 4°C on a shaker. After washing with PBST, the samples were incubated with secondary antibodies conjugated to Alexa Fluor (1:500) and Phalloidin conjugated to Alexa Fluor (1:500) for 1 hour at RT on a shaker. Samples were then washed with PBST, mounted on glass slides with an aqueous mounting medium, and, for fish neuromast samples, underwent an additional fixation before being equilibrated with glycerol.

### Explant Culture

BPs were cultured using a 3D-collagen droplet method as previously described (Singh et al 2022). Briefly, a mixture of rat tail collagen (400µl), 10XDMEM (50µl), 7.5% Sodium Bicarbonate (30µl), and HEPES (10µl) was used to form droplets in 4-well plates. One cochlea was placed in each drop, which was then incubated for 10 minutes at 37°C. Afterwards, culture media (DMEM+N2), with or without 10 µM Y-27632 ROCK inhibitor, was added.

### Live Imaging

We used a previously described method to perform live imaging (Prakash, Weninger et al 2025). In brief, we dissected BP in ice-cold HBSS and mounted it using a collagen droplet on a coverslipped 35 mm Petri dish. We then incubated the BP with 50nM Sir-Actin for 30mins at 37°C incubator. Excess SIR-Actin was washed off by rinsing it twice with HBSS. We placed the 35mm dish in a 37°C incubator with 5%CO_2_ on an inverted microscope with a 60× oil immersion objective with NA 1.42 and imaged using a Cy5 filter, taking images every 5 minutes.

### Imaging

All images are acquired using an Olympus FV3000 system equipped with a 60X oil immersion objective with 1.42 NA, operated by Olympus Fluoview software. Images are acquired at a Nyquist sampling rate with a pinhole size of 0.8-1 airy disk and a step size of 0.5 μm.

### Image-Analysis

To analyze the geometry and morphology of epithelium, we segmented images using the Tissue-analyser plugin in FIJI. We then transferred data from HCA or beta-spectrin II staining, along with the presence of stereocilia indicating HCs, to identify HCs and record their X-Y coordinates as CSV files. Using this CSV file and the segmented images as input for a custom MATLAB script, we calculated parameters such as apical surface area, perimeter, neighbor count, and circularity for each cell, distinguishing between HC and SC.

For relative intensity assessments, lines of interest about 3 pixels thick were drawn at HC-SC and SC-SC junctions, and the mean intensity along these lines was measured. The resulting intensity values were normalized to the maximum intensity for each image and plotted with GraphPad Prism software.

To evaluate fluctuations in HC and SC shape, we segmented each frame of live-imaging videos to calculate changes in circularity for HCs and SCs over hourly intervals.

**Movie 1: Live imaging of control E10 BP.**

E10 BP stained with SiR-Actin (50nM, 30 mins) and imaged in explant for 5 hours with images taken after every 5 minutes showing HCs to maintain its shape while SCs to change shape dynamically.

**Movie 2: Live imaging of Y-27632 treated E10 BP.**

E10 BP stained with SiR-Actin (50nM, 30 mins) and treated with Y-27632 (10µM) during imaging of explant for 5 hours with images taken after every 5 minutes.

## Supplementary Figures

**Figure S1.**
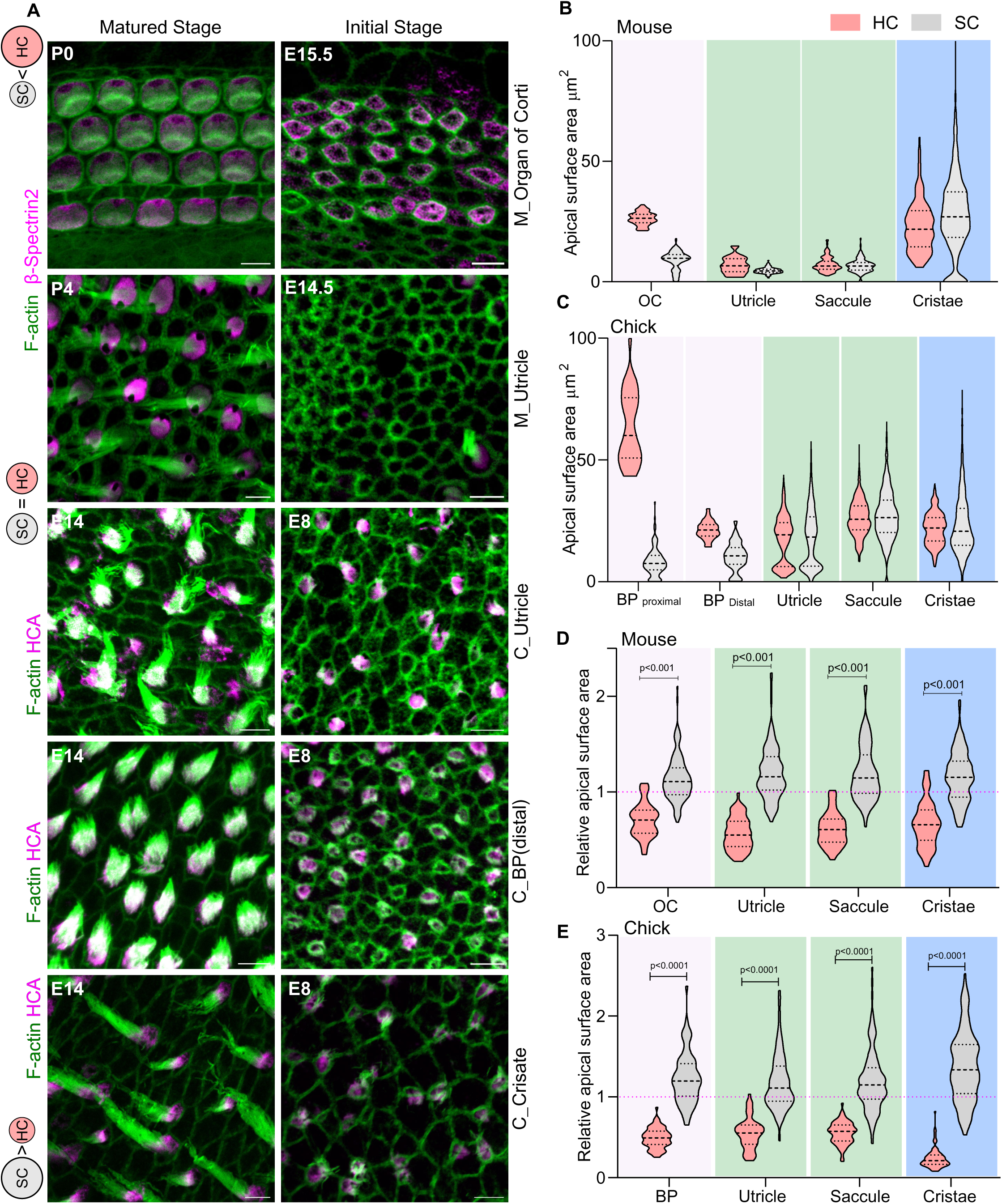
Across epithelia, HCs are smaller than SCs at the initial stages that diverge into distinct areas in development. A. Sensory epithelia of mouse Cristae, saccule and chick BP and saccule at the matured stage (P4 for mouse and E14 for chick) and initial stage (E14.5 for mouse and E8 for chick) stained for F-actin (green) and hair cell marker (HCA, Beta Spectrin 2). B. Apical Surface area of HC and SC from matured mouse sensory epithelia. C. Apical surface area of HCs and SCs from inner ear sensory epithelia of chick at E14. D. Relative apical surface area of HCs and SCs from mouse inner ear sensory epithelia at E14.5. E. Relative apical surface area of HCs and SCs from chick inner ear sensory epithelia at E8.

**Figure S2.**
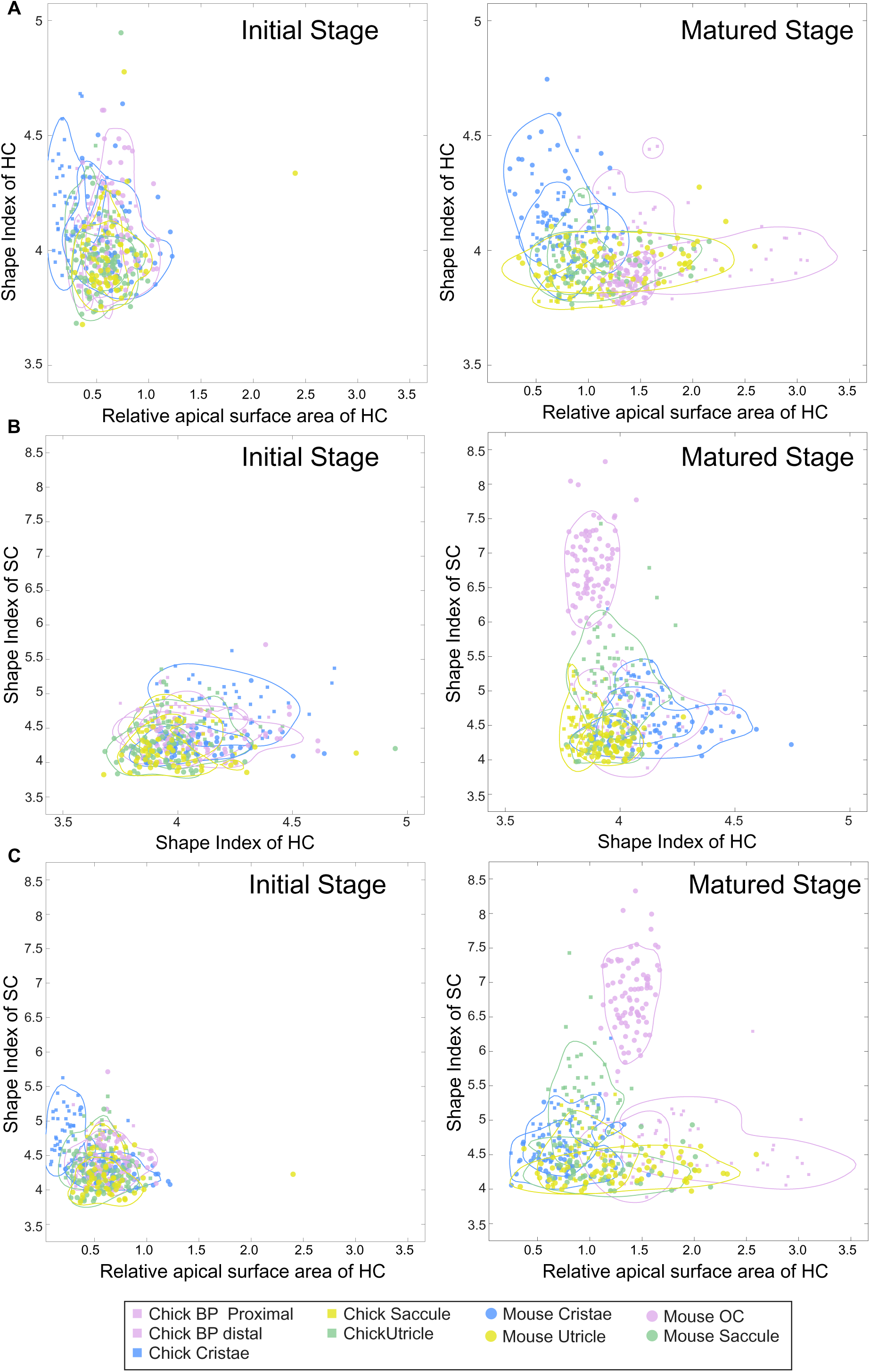
HCs are smaller and circular compared to SCs at initiation. A. Two-feature plots for the relative apical surface area and shape index of HC of inner ear sensory epithelia from mouse and chick. B. Two-feature plots for the shape index of SC and shape index of HC of inner ear sensory epithelia from mouse and chick. C. Two-feature plots for the relative apical surface area and shape index of SC of inner ear sensory epithelia from mouse and chick.

**Figure S3.**
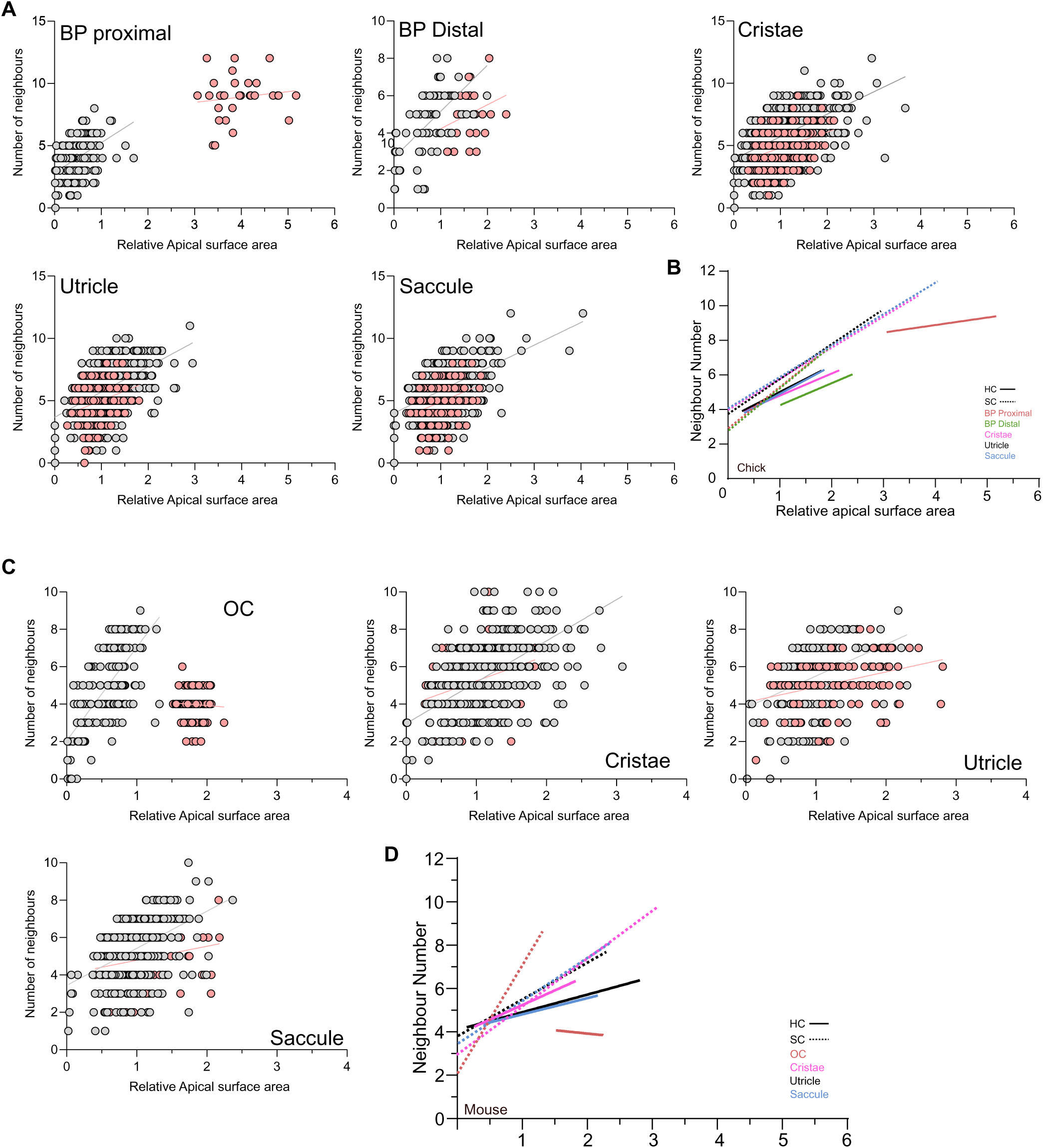
HCs and SCs show a distinct relationship between area and neighbour number. A. Number of neighbours with respect to relative apical surface area for HC and SC in chick sensory epithelia. B. Regression line representing the number of neighbours with respect to relative apical surface area for HC and SC in chick sensory epithelia. C. Number of neighbours with respect to relative apical surface area for HC and SC in Mouse sensory epithelia. D. Regression line representing the number of neighbours with respect to relative apical surface area for HC and SC in chick sensory epithelia.

**Figure S4.**
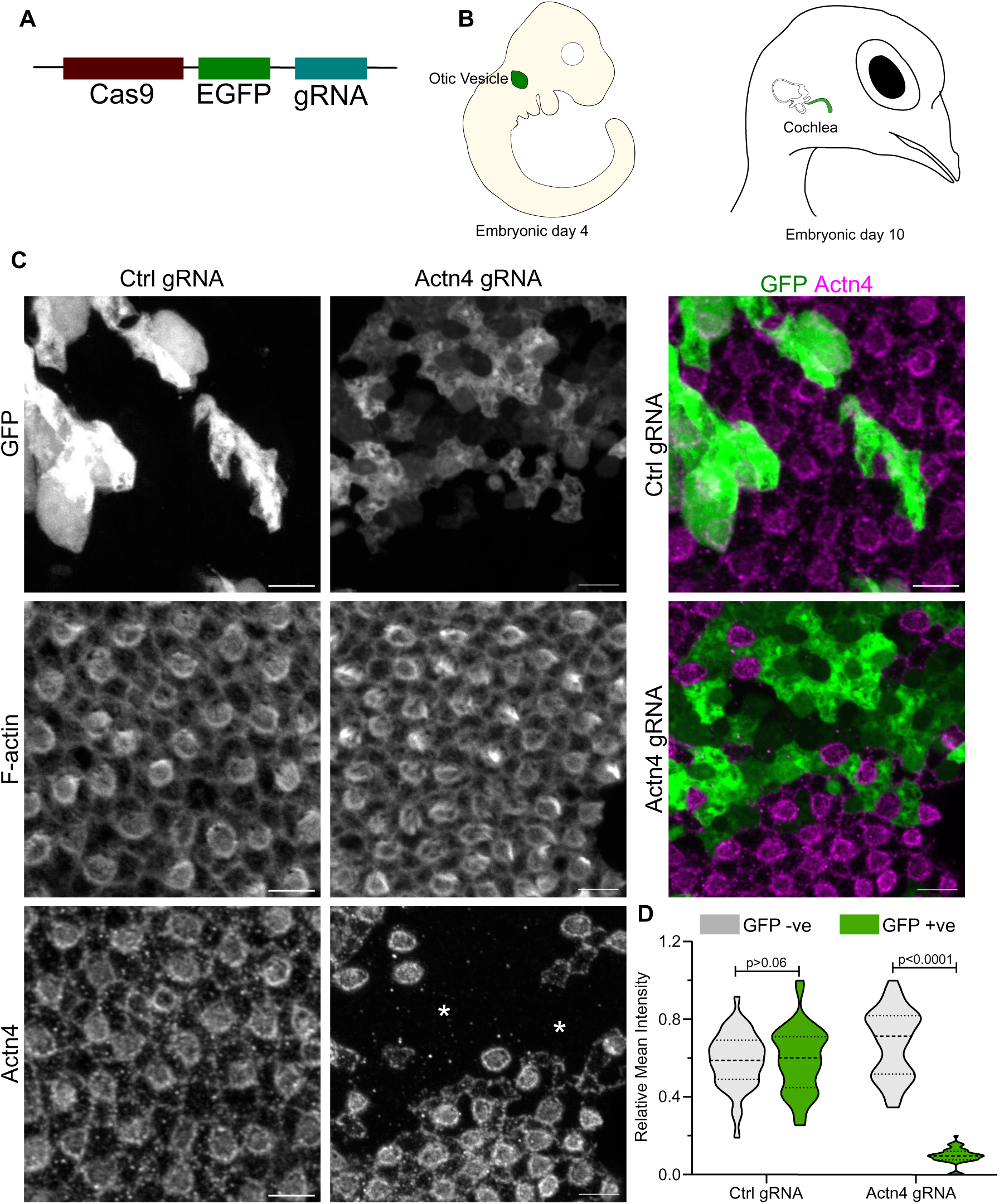
Electroporation of gRNA targetting Actn4 mosaically knockouts Actn4. A. Design of DNA construct for electroporation. B. Schematic of electroporation in primordia of inner ear (otic vesicle) at E3.5 and grown till E10. C. E10 BP electroporated with ctrl gRNA, Actn4 gRNA stained for F-actin (grey), Actn4 (grey, magenta) and GFP (grey, green). D. Relative mean intensity of Actn4 in GFP negative and positive patches from Ctrl and Actn4 gRNA electroporated patches. N=4 samples. Scale Bar: 5µm Statistics: Unpaired-Test

**Figure S5.**
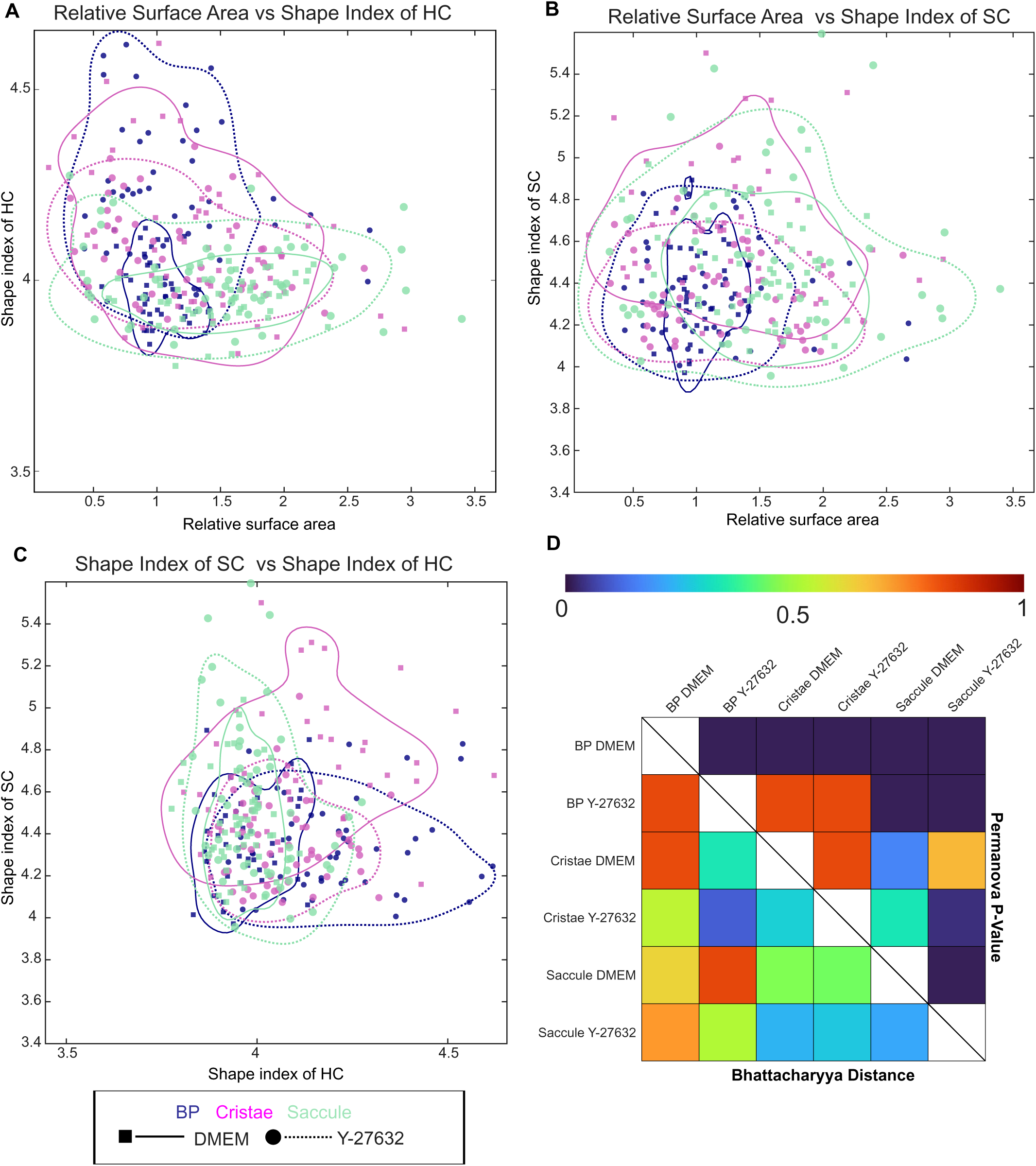
Selective accumulation of adhesion and contractility proteins on junctions. A. Two-feature plots for the relative apical surface area and shape index of HC of control (DMEM) and myosin phosphorylation inhibitor (Y-27632) treated sensory epithelia. B. Two-feature plots for the shape index of SC and shape index of HC of control (DMEM) and myosin phosphorylation inhibitor (Y-27632) treated sensory epithelia. C. Two-feature plots for the relative apical surface area and shape index of control (DMEM) and myosin phosphorylation inhibitor (Y-27632) treated sensory epithelia. D. Bhattacharyya distance and the Permanova statistical test to quantify the overlap and distinction between the control (DMEM) and myosin phosphorylation inhibitor (Y-27632) treated sensory epithelia.

## References

Andrews, T.G.R. et al. (2021) “Single-cell morphometrics reveals ancestral principles of notochord development,” Development, 148(16), p. dev199430. Available at: 10.1242/dev.199430.

Bailles, A., Gehrels, E.W. and Lecuit, T. (2022) “Mechanochemical Principles of Spatial and Temporal Patterns in Cells and Tissues,” Annual Review of Cell and Developmental Biology, 38(1), pp. 321–347. Available at: 10.1146/annurev-cellbio-120420-095337.

Basch, M.L. et al. (2016) “Where hearing starts: the development of the mammalian cochlea,” Journal of Anatomy, 228(2), pp. 233–254. Available at: 10.1111/joa.12314.

Battistara, M. et al. (2026) “Morphospace analysis reveals divergent cellular behaviours driving tissue internalisation during insect gastrulation.” Available at: 10.64898/2026.06.30.735526.

Bermingham, N.A. et al. (1999) “Math1: an essential gene for the generation of inner ear hair cells,” *Science (New York*, N.Y.), 284(5421), pp. 1837–1841. Available at: 10.1126/science.284.5421.1837.

Budd, G.E. (2021) “Morphospace,” Current Biology, 31(19), pp. R1181–R1185. Available at: 10.1016/j.cub.2021.08.040.

Burns, J.C. et al. (2012) “Over half the hair cells in the mouse utricle first appear after birth, with significant numbers originating from early postnatal mitotic production in peripheral and striolar growth zones,” Journal of the Association for Research in Otolaryngology: JARO, 13(5), pp. 609–627. Available at: 10.1007/s10162-012-0337-0.

Burns, J.C. et al. (2015) “Single-cell RNA-Seq resolves cellular complexity in sensory organs from the neonatal inner ear,” Nature Communications, 6, p. 8557. Available at: 10.1038/ncomms9557.

Burns, J.C. and Stone, J.S. (2017) “Development and regeneration of vestibular hair cells in mammals,” Seminars in Cell & Developmental Biology, 65, pp. 96–105. Available at: 10.1016/j.semcdb.2016.11.001.

Cohen, R. et al. (2020) “Mechanical forces drive ordered patterning of hair cells in the mammalian inner ear,” Nature Communications, 11(1), p. 5137. Available at: 10.1038/s41467-020-18894-8.

Collinet, C. and Lecuit, T. (2021) “Programmed and self-organized flow of information during morphogenesis,” Nature Reviews Molecular Cell Biology, 22(4), pp. 245–265. Available at: 10.1038/s41580-020-00318-6.

Curran, S. et al. (2017) “Myosin II Controls Junction Fluctuations to Guide Epithelial Tissue Ordering,” Developmental Cell, 43(4), pp. 480–492.e6. Available at: 10.1016/j.devcel.2017.09.018.

Driver, E.C. et al. (2008) “Hedgehog Signaling Regulates Sensory Cell Formation and Auditory Function in Mice and Humans,” The Journal of Neuroscience, 28(29), pp. 7350–7358. Available at: 10.1523/JNEUROSCI.0312-08.2008.

Duguay, D., Foty, R.A. and Steinberg, M.S. (2003) “Cadherin-mediated cell adhesion and tissue segregation: qualitative and quantitative determinants,” Developmental Biology, 253(2), pp. 309–323. Available at: 10.1016/s0012-1606(02)00016-7.

Ehrlicher, A.J. et al. (2015) “Alpha-actinin binding kinetics modulate cellular dynamics and force generation,” Proceedings of the National Academy of Sciences of the United States of America, 112(21), pp. 6619–6624. Available at: 10.1073/pnas.1505652112.

Farhadifar, R. et al. (2007) “The influence of cell mechanics, cell-cell interactions, and proliferation on epithelial packing,” Current biology: CB, 17(24), pp. 2095–2104. Available at: 10.1016/j.cub.2007.11.049.

Gnedeva, K. et al. (2017) “Elastic force restricts growth of the murine utricle,” eLife, 6, p. e25681. Available at: 10.7554/eLife.25681.

Goodyear, R. and Richardson, G. (1997) “Pattern formation in the basilar papilla: evidence for cell rearrangement,” The Journal of Neuroscience: The Official Journal of the Society for Neuroscience, 17(16), pp. 6289–6301. Available at: 10.1523/JNEUROSCI.17-16-06289.1997.

Groves, A.K. and Fekete, D.M. (2012) “Shaping sound in space: the regulation of inner ear patterning,” *Development (Cambridge*, England), 139(2), pp. 245–257. Available at: 10.1242/dev.067074.

Heisenberg, C.-P. and Bellaïche, Y. (2013) “Forces in Tissue Morphogenesis and Patterning,” Cell, 153(5), pp. 948–962. Available at: 10.1016/j.cell.2013.05.008.

Huh, S.-H., Warchol, M.E. and Ornitz, D.M. (2015) “Cochlear progenitor number is controlled through mesenchymal FGF receptor signaling,” eLife, 4, p. e05921. Available at: 10.7554/eLife.05921.

Hunter, G.L. et al. (2016) “Coordinated control of Notch-Delta signalling and cell cycle progression drives lateral inhibition mediated tissue patterning,” *Development*, p. dev.134213. Available at: 10.1242/dev.134213.

Jolly, M.K. et al. (2015) “Operating principles of Notch–Delta–Jagged module of cell–cell communication,” New Journal of Physics, 17(5), p. 055021. Available at: 10.1088/1367-2630/17/5/055021.

Kailath, T. (1967) “The Divergence and Bhattacharyya Distance Measures in Signal Selection,” IEEE Transactions on Communications, 15(1), pp. 52–60. Available at: 10.1109/TCOM.1967.1089532.

Kasirer, S. and Sprinzak, D. (2024) “Interplay between Notch signaling and mechanical forces during developmental patterning processes,” Current Opinion in Cell Biology, 91, p. 102444. Available at: 10.1016/j.ceb.2024.102444.

Katsunuma, S. et al. (2016) “Synergistic action of nectins and cadherins generates the mosaic cellular pattern of the olfactory epithelium,” Journal of Cell Biology, 212(5), pp. 561–575. Available at: 10.1083/jcb.201509020.

Kemp, J.P. and Brieher, W.M. (2018) “The actin filament bundling protein α-actinin-4 actually suppresses actin stress fibers by permitting actin turnover,” Journal of Biological Chemistry, 293(37), pp. 14520–14533. Available at: 10.1074/jbc.RA118.004345.

Khalaily, L. et al. (2026) “Live imaging and multimodal profiling reveal transdifferentiation of a cochlear supporting cell subpopulation upon Notch inhibition,” Science Advances, 12(25), p. eaed3887. Available at: 10.1126/sciadv.aed3887.

Kokic, M. et al. (2019) “Minimisation of surface energy drives apical epithelial organisation and gives rise to Lewis’ law.” Biophysics. Available at: 10.1101/590729.

Lanford, P.J. et al. (1999) “Notch signalling pathway mediates hair cell development in mammalian cochlea,” Nature Genetics, 21(3), pp. 289–292. Available at: 10.1038/6804.

Lewis, F.T. (1926) “The effect of cell division on the shape and size of hexagonal cells,” The Anatomical Record, 33(5), pp. 331–355. Available at: 10.1002/ar.1090330502.

Li, X.-J. et al. (2025) “The Notch ligand Jagged1 plays a dual role in cochlear hair cell regeneration,” Nature Communications, 16(1), p. 8169. Available at: 10.1038/s41467-025-63053-6.

Lush, M.E. et al. (2019) “scRNA-Seq reveals distinct stem cell populations that drive hair cell regeneration after loss of Fgf and Notch signaling,” eLife, 8, p. e44431. Available at: 10.7554/eLife.44431.

Mann, Z.F. et al. (2014) “A gradient of Bmp7 specifies the tonotopic axis in the developing inner ear,” Nature Communications, 5, p. 3839. Available at: 10.1038/ncomms4839.

Mann, Z.F. et al. (2017) “Shaping of inner ear sensory organs through antagonistic interactions between Notch signalling and Lmx1a,” eLife, 6, p. e33323. Available at: 10.7554/eLife.33323.

McKenzie, E., Krupin, A. and Kelley, M.W. (2004) “Cellular growth and rearrangement during the development of the mammalian organ of Corti,” Developmental Dynamics, 229(4), pp. 802–812. Available at: 10.1002/dvdy.10500.

Mendieta-Serrano, M.A. et al. (2025) “A structural transition ensures robust formation of skeletal muscle.” Developmental Biology. Available at: 10.1101/2025.09.22.677369.

Muthukrishnan, S. et al. (2026) “Glassy dynamics in active epithelia emerge from an interplay of mechanochemical feedback and crowding,” Nature Communications [Preprint]. Available at: 10.1038/s41467-026-74163-0.

Nommick, A. et al. (2025) “Dual role of Xenopus Odf2 in multiciliated cell patterning and differentiation,” Developmental Biology, 520, pp. 224–238. Available at: 10.1016/j.ydbio.2025.01.014.

Ono, K., et al. (2014) “FGFR1-Frs2/3 Signalling Maintains Sensory Progenitors during Inner Ear Hair Cell Formation,” PLoS Genetics. Edited by K.S.E. Cheah, 10(1), p. e1004118. Available at: 10.1371/journal.pgen.1004118.

Pie, M.R. and Weitz, J.S. (2005) “A Null Model of Morphospace Occupation,” The American Naturalist, 166(1), pp. E1–E13. Available at: 10.1086/430727.

Prakash, A., Raman, S., et al. (2025) “Coupling between spatial compartments integrates morphogenetic patterning in the organ of Corti,” PLOS Biology. Edited by A.G. Cheng, 23(9), p. e3003350. Available at: 10.1371/journal.pbio.3003350.

Prakash, A., Weninger, J., et al. (2025) “Junctional force patterning drives both positional order and planar polarity in the auditory epithelia,” Nature Communications, 16(1), p. 3927. Available at: 10.1038/s41467-025-58557-0.

Puligilla, C. et al. (2007) “Disruption of fibroblast growth factor receptor 3 signaling results in defects in cellular differentiation, neuronal patterning, and hearing impairment,” Developmental Dynamics: An Official Publication of the American Association of Anatomists, 236(7), pp. 1905–1917. Available at: 10.1002/dvdy.21192.

Rombouts, J., Elliott, J. and Erzberger, A. (2023) “Forceful patterning: theoretical principles of mechanochemical pattern formation,” The EMBO Reports, 24(12), p. EMBR202357739. Available at: 10.15252/embr.202357739.

Salbreux, G. et al. (2012) “Coupling mechanical deformations and planar cell polarity to create regular patterns in the zebrafish retina,” PLoS computational biology, 8(8), p. e1002618. Available at: 10.1371/journal.pcbi.1002618.

Shaya, O. et al. (2017) “Cell-Cell Contact Area Affects Notch Signaling and Notch-Dependent Patterning,” Developmental Cell, 40(5), pp. 505–511.e6. Available at: 10.1016/j.devcel.2017.02.009.

Singh, N. et al. (2022) “In Ovo and Ex Ovo Methods to Study Avian Inner Ear Development,” Journal of Visualized Experiments, (184), p. 64172. Available at: 10.3791/64172.

Singh, N. et al. (2024) “Mosaic Atoh1 deletion in the chick auditory epithelium reveals a homeostatic mechanism to restore hair cell number,” Developmental Biology, 516, pp. 35–46. Available at: 10.1016/j.ydbio.2024.07.017.

Thiede, B.R. et al. (2014) “Retinoic acid signalling regulates the development of tonotopically patterned hair cells in the chicken cochlea,” Nature Communications, 5, p. 3840. Available at: 10.1038/ncomms4840.

Togashi, H. et al. (2011) “Nectins establish a checkerboard-like cellular pattern in the auditory epithelium,” *Science (New York*, N.Y.), 333(6046), pp. 1144–1147. Available at: 10.1126/science.1208467.

Tsai, T.Y.-C. et al. (2020) “An adhesion code ensures robust pattern formation during tissue morphogenesis,” *Science (New York*, N.Y.), 370(6512), pp. 113–116. Available at: 10.1126/science.aba6637.

Wang, X. et al. (2020) “Anisotropy links cell shapes to tissue flow during convergent extension,” Proceedings of the National Academy of Sciences of the United States of America, 117(24), pp. 13541–13551. Available at: 10.1073/pnas.1916418117.

Weninger, J. et al. (2026) “Force patterning drives quasistratification and graded tissue-scale spatial order in auditory epithelia,” Proceedings of the National Academy of Sciences, 123(19), p. e2519341123. Available at: 10.1073/pnas.2519341123.

Wibowo, I. et al. (2011) “Compartmentalized Notch signaling sustains epithelial mirror symmetry,” *Development (Cambridge*, England), 138(6), pp. 1143–1152. Available at: 10.1242/dev.060566.

Wickström, S.A. and Niessen, C.M. (2018) “Cell adhesion and mechanics as drivers of tissue organization and differentiation: local cues for large scale organization,” Current Opinion in Cell Biology, 54, pp. 89–97. Available at: 10.1016/j.ceb.2018.05.003.

Yang, X. et al. (2017) “Establishment of planar cell polarity is coupled to regional cell cycle exit and cell differentiation in the mouse utricle,” Scientific Reports, 7(1), p. 43021. Available at: 10.1038/srep43021.

Zaidel-Bar, R. and Agarwal, P. (2025) “Outside influences: The impact of extracellular matrix mechanics on cell migration,” Current Topics in Developmental Biology. Elsevier, pp. 29–65. Available at: 10.1016/bs.ctdb.2025.01.003.

