## Supplementary material for "Coexistence of asymmetric cell shape dynamics governs organisational diversity in sensory epithelia": Code used for analysis

```
cd 'E:\Appy\Neighbour_number\' %%%%% Destination
number_files = 54; %%%%%%%%%%%%%% Number of Files
for kk = 1:number_files
File_Name = strcat('N',string(kk),'.tif')
mat = strcat('N',string(kk),'.csv')
Og_file = imread(File_Name);
Image_file = imcomplement(logical(Og_file(:,:,1)));
[x,y] = size(Image_file);
Image_file([1,x],:)= [];
Image_file(:,[1,y]) = [];
Image_file_true = imclearborder(Image_file,4);
Connected_comp = bwconncomp(Image_file_true,4);



dim = 2;
[xx,yy] = ndgrid(-dim:dim);
nhood = sqrt(xx.^2 + yy.^2 ) <= dim;
Props =
regionprops(Connected_comp,'Centroid','Area','PixelIdxList',"Circularity","Solidity","MaxFeretPro
perties","Orientation","ConvexHull","BoundingBox");
imshow(Image_file_true)
hold on
for k = 1:size(Props,1)
   text(Props(k).Centroid(1),Props(k).Centroid(2), ...
   sprintf('%1.f', k), ...
   'Color','r');
end
hold off
Centroid_All = vertcat(Props.Centroid);
%ID_Cell = zeros(20,size(Props,1));
Count = zeros(1,size(Props,1));
selected_Cell = readmatrix(mat);
selected_Cell_Coordinates = selected_Cell(:,[4,5]);
selected_Cell_Coordinates(:,1) = ceil(selected_Cell_Coordinates(:,1));
selected_Cell_Coordinates(:,2) = ceil(selected_Cell_Coordinates(:,2));
Select_Mask = zeros(size(Og_file(:,:,1)));
for k = 2:size(selected_Cell_Coordinates,1)
Select_Mask(selected_Cell_Coordinates(k,2),selected_Cell_Coordinates(k,1))=1;
end
%%%%%%%%%%%%%%%%%%%%%%%%%%%%%%%%%%%%%%%%%%%
%%%%%%%%%%%%%%%%%%%%%%%%%%%%%%%%%%%%%%%%%%%
%%%%%%%%%%%%%%%%%%%%%%%%%%%%%%%%%%%%%%%%%%%
%%%%%%%%%%%%%%%%%%%%%%%%%%%%%%%%%%%%%%
Select_Mask([1,x],:)= [];
Select_Mask(:,[1,y]) = [];
Cell_A =  imreconstruct(logical(Select_Mask),Image_file_true,4);
Connected_comp_A = bwconncomp(Cell_A,4);
Cell_A_Props =
regionprops(Connected_comp_A,'Centroid','Area','PixelIdxList',"Circularity","Solidity","MaxFeret
Properties","Orientation","ConvexHull","BoundingBox","Perimeter");
Cell_B = Image_file_true-Cell_A;
Connected_comp_B = bwconncomp(Cell_B,4);
Cell_B_Props =
regionprops(Connected_comp_B,'Centroid','Area','PixelIdxList',"Circularity","Solidity","MaxFeret
Properties","Orientation","ConvexHull","BoundingBox","Perimeter");
%%%%%%%%%%%%%%%%%%%%%%%%%%%%%%%%%%%%%%%%%%%
%%%%%%%%%%%%%%%%%%%%%%%%%%%%%%%%%%%%%%%%%%%
%%%%%%%%%%%%%%%%%%%%%%%%%%%%%%%%%%%%%%%%%%%
%%%%%%%%%%%%%%%%%%%%%%%%%%%%%%%%%%%%%%%
```

```
Prop_A_Neighbour = zeros(1,size(Cell_A_Props,1));
CellA_A_Neighbour = zeros(1,size(Cell_A_Props,1));
CellA_B_Neighbour = zeros(1,size(Cell_A_Props,1));
Cell_A = logical(Cell_A);
Cell_B = logical(Cell_B);
for k = 1:size(Cell_A_Props,1)
    Mask = zeros(size(Image_file_true));
    Mask(Cell_A_Props(k).PixelIdxList)=1;
    Dilate_Image = imdilate(Mask,nhood);
    Boundary_Image = Dilate_Image - Mask;
    Boundary_Image = logical(Boundary_Image);
    J = imreconstruct(Boundary_Image,Image_file_true,4);
    new_rgb = cat(3,Mask,J,zeros(size(Image_file_true)));
    Cell_rgb = sprintf('Cell_type_A_%1.f.tiff', k)
    Neighbour_comp = bwconncomp(J,4);
    Neighbour_Props =
regionprops(Neighbour_comp,'Centroid','Area','PixelIdxList',"Circularity","Solidity","MaxFeretPro
perties","Orientation","ConvexHull","BoundingBox");
    Prop_A_Neighbour(k) = size(Neighbour_Props,1);
    %%%%%%%%%%%%%%%%%%%%%%%%%%%%%%%%%%%%%%%%%%%%%
%%%%%%%%%%%%%%%%%%%%%%%%%%%%%%%%%%%%%%%%%%%%%%
%%%%%%%%%%%%%%%%%%%%%%%%%%%%%%%%%%%%%%%%%55555
    JAA = imreconstruct(Boundary_Image,Cell_A,4);
    Neighbour_comp_AA = bwconncomp(JAA,4);
    Neighbour_Props_AA =
regionprops(Neighbour_comp_AA,'Centroid','Area','PixelIdxList',"Circularity","Solidity","MaxFeret
Properties","Orientation","ConvexHull","BoundingBox");
    CellA_A_Neighbour(k) = size(Neighbour_Props_AA,1);
    %%%%%%%%%%%%%%%%%%%%%%%%%%%%%%%%%%%%%%%%%%%%%
%%%%%%%%%%%%%%%%%%%%%%%%%%%%%%%%%%%%%%%%%%%%%%
%%%%%%%%%%%%%%%%%%%%%%%%%%%%%%%%%%%%%%%%%%%%%%
%%%%%%%%%%5
    JAB = imreconstruct(Boundary_Image,Cell_B,4);
    Neighbour_comp_AB = bwconncomp(JAB,4);
    Neighbour_Props_AB =
regionprops(Neighbour_comp_AB,'Centroid','Area','PixelIdxList',"Circularity","Solidity","MaxFeret
Properties","Orientation","ConvexHull","BoundingBox");
    CellA_B_Neighbour(k) = size(Neighbour_Props_AB,1);
    %%%%%%%%%%%%%%%%%%%%%%%%%%%%%%%%%%%%%%%%%%%%%
%%%%%%%%%%%%%%%%%%%%%%%%%%%%%%%%%%%%%%%%%%%%%%
%%%%%%%%%%%%%%%%%%%%%%%%%%%%%%%%%%%%%%%%%%%%%%
%%%%%%%%%%%%%%%%%%%

    Centroid_Neighbour = vertcat(Neighbour_Props.Centroid);
    indices = find(ismember(Centroid_All, Centroid_Neighbour,'rows'));
    ID_Cell{k}= indices;
    Count(k) = size(indices,1);
    Location{k} = Props(k).PixelIdxList;
end
table_A = struct2table(Cell_A_Props);
table_A.("Neighbour Number") = Prop_A_Neighbour';
table_A.("A Neighbour Number") = CellA_A_Neighbour';
table_A.("B Neighbour Number") = CellA_B_Neighbour';

table_A.ConvexHull = [];
table_A.PixelIdxList = [];

Prop_B_Neighbour = zeros(1,size(Cell_B_Props,1));
CellB_A_Neighbour = zeros(1,size(Cell_A_Props,1));
CellB_B_Neighbour = zeros(1,size(Cell_A_Props,1));
```

```
for k = 1:size(Cell_B_Props,1)
    Mask = zeros(size(Image_file_true));
    Mask(Cell_B_Props(k).PixelIdxList)=1;
    Dilate_Image = imdilate(Mask,nhood);
    Boundary_Image = Dilate_Image - Mask;
    Boundary_Image = logical(Boundary_Image);
    J = imreconstruct(Boundary_Image,Image_file_true,4);
    new_rgb = cat(3,Mask,J,zeros(size(Image_file_true)));
    Cell_rgb = sprintf('Cell_type_B_%1.f.tiff', k)

    Neighbour_comp = bwconncomp(J,4);
    Neighbour_Props =
regionprops(Neighbour_comp,'Centroid','Area','PixelIdxList',"Circularity","Solidity","MaxFeretPro
perties","Orientation","ConvexHull","BoundingBox");
    Prop_B_Neighbour(k) = size(Neighbour_Props,1);
    %%%%%%%%%%%%%%%%%%%%%%%%%%%%%%%%%%%%%%%%%%%%
%%%%%%%%%%%%%%%%%%%%%%%%%%%%%%%%%%%%%%%%%%%%%
%%%%%%%%%%%%%%%%%%%%%%%%%%%%%%%%%%%%%%%%55555
    JBA = imreconstruct(Boundary_Image,Cell_A,4);
    Neighbour_comp_BA = bwconncomp(JBA,4);
    Neighbour_Props_BA =
regionprops(Neighbour_comp_BA,'Centroid','Area','PixelIdxList',"Circularity","Solidity","MaxFeret
Properties","Orientation","ConvexHull","BoundingBox");
    CellB_A_Neighbour(k) = size(Neighbour_Props_BA,1);
    %%%%%%%%%%%%%%%%%%%%%%%%%%%%%%%%%%%%%%%%%%%%
%%%%%%%%%%%%%%%%%%%%%%%%%%%%%%%%%%%%%%%%%%%%%
%%%%%%%%%%%%%%%%%%%%%%%%%%%%%%%%%%%%%%%%%%%%%
%%%%%%%%%%5
    JBB = imreconstruct(Boundary_Image,Cell_B,4);
    Neighbour_comp_BB = bwconncomp(JBB,4);
    Neighbour_Props_BB =
regionprops(Neighbour_comp_BB,'Centroid','Area','PixelIdxList',"Circularity","Solidity","MaxFeret
Properties","Orientation","ConvexHull","BoundingBox");
    CellB_B_Neighbour(k) = size(Neighbour_Props_BB,1);
    %%%%%%%%%%%%%%%%%%%%%%%%%%%%%%%%%%%%%%%%%%%%
%%%%%%%%%%%%%%%%%%%%%%%%%%%%%%%%%%%%%%%%%%%%%
%%%%%%%%%%%%%%%%%%%%%%%%%%%%%%%%%%%%%%%%%%%%%
%%%%%%%%%%%%%%%%%%%
    Centroid_Neighbour = vertcat(Neighbour_Props.Centroid);
    indices = find(ismember(Centroid_All, Centroid_Neighbour,'rows'));
    ID_Cell{k}= indices;
    Count(k) = size(indices,1);
    Location{k} = Props(k).PixelIdxList;
end

table_B = struct2table(Cell_B_Props);
table_B.("Neighbour Number") = Prop_B_Neighbour';
table_B.("A Neighbour Number") = CellB_A_Neighbour';
table_B.("B Neighbour Number") = CellB_B_Neighbour';

table_B.ConvexHull = [];
table_B.PixelIdxList = [];
writetable(table_A,'Cell_Type_A_Props.xlsx');
writetable(table_B,'Cell_Type_B_Props.xlsx');
end
```
